# A dynamic mechanism for allosteric activation of Aurora kinase A by activation loop phosphorylation

**DOI:** 10.1101/205260

**Authors:** Emily F. Ruff, Joseph M. Muretta, Andrew Thompson, Eric Lake, Soreen Cyphers, Steven K. Albanese, Sonya M. Hanson, Julie M. Behr, David D. Thomas, John D. Chodera, Nicholas M. Levinson

**Author notes:** present address: Department of Chemistry, Winona State University, Winona, Minnesota, USA.

## Abstract

Many eukaryotic protein kinases are activated by phosphorylation on a specific conserved residue in the regulatory activation loop, a post-translational modification thought to stabilize the active DFG-In state of the catalytic domain. Here we use a battery of spectroscopic methods that track different catalytic elements of the kinase domain to show that the ~100-fold activation of the mitotic kinase Aurora A (AurA) by phosphorylation occurs without a population shift to the DFG-In state, and that the activation loop of the activated kinase remains highly dynamic. Instead, molecular dynamics simulations and electron paramagnetic resonance experiments show that phosphorylation profoundly alters the structure and dynamics of the DFG-In subpopulation, leading to activation of the kinase. Kinetics experiments tracking structural transitions during nucleotide binding suggest that a substantial DFG-Out subpopulation is an important feature of activated AurA that evolved to optimize the kinetics of substrate binding and product release.

## Introduction

Protein phosphorylation is a central feature of cellular signal transduction, and stringent regulatory control of the participating protein kinases is critical for the integrity of these pathways. Kinase activity is typically regulated by finely-tuned allosteric mechanisms that reversibly switch the kinase domain between active and inactive conformational states^1^. Disruption of these mechanisms, leading to constitutive kinase activity, is a major cause of cancer, and small molecules that inhibit specific disease-associated kinases constitute an important class of modern cancer drugs^2^.

Phosphorylation on a specific site in the activation loop of the kinase domain is the most widely conserved regulation mechanism in protein kinases^3^. X-ray structures suggest that ionic interactions between the phosphate moiety and a pocket of basic residues lock the activation loop into a conserved active conformation^4–6^. In this conformation, the catalytic asp-phe-gly (DFG) motif at the N-terminal end of the activation loop adopts an active “DFG-In” conformation, in which the aspartate residue of the DFG motif points into the active site to coordinate Mg-ATP.

In the absence of phosphorylation protein kinases often adopt autoinhibited states, in which activity is blocked by conformational rearrangements of the activation loop and DFG motif. An important mode of autoinhibition involves a flip of the aspartate residue of the DFG motif out of the active site, preventing magnesium coordination^7–9^. Many protein kinases have been observed to adopt these “DFG-Out” states, and some have been targeted with small-molecule inhibitors that preferentially bind to the DFG-Out conformation^10^. In addition to phosphorylation, kinase conformation is typically also modulated by the binding of accessory proteins that further tune the activity level of the enzyme.

The serine/threonine kinase Aurora A (AurA) is an essential mitotic protein that controls many cellular processes including mitotic spindle assembly, centrosome maturation, and mitotic entry^11–15^. These functions of AurA are driven by two distinct activation mechanisms of the kinase operating in different spatiotemporal contexts. At the centrosome, AurA is activated by autophosphorylation on the activation loop residue T288. In contrast, at the mitotic spindle, AurA is activated by the spindle-associated protein Tpx2^16^, and this pool of the kinase is kept in the unphosphorylated state by the continual action of the phosphatase PP6^17,18^.

Extensive *in vitro* studies have shown that Tpx2 and phosphorylation act independently to increase AurA kinase activity by up to several hundred-fold^19,20^. Binding of Tpx2 to unphosphorylated AurA triggers a population shift from a DFG-Out to the DFG-In state in solution^21^. Crystal structures of phosphorylated AurA bound to Tpx2 show that the T288 phosphothreonine residue forms extensive ionic interactions unique to the DFG-In state, suggesting that both factors simply stabilize the same active conformation^22–24^. In this paper, we show that phosphorylation on T288 in fact activates AurA through a completely different mechanism than Tpx2. Using three complementary spectroscopic methods we show that phosphorylation does not trigger a switch to the DFG-In state, and that the phosphorylated activation loop of AurA continually samples both active and inactive conformational states. Instead, phosphorylation acts by enhancing the catalytic activity of the subpopulation of molecules adopting the DFG-In state. Stopped flow kinetics experiments point to the DFG-Out state being important for nucleotide dissociation, and suggest that the transition between DFG-Out and DFG-In states may be a central feature of the catalytic cycle of the enzyme.

## Results and Discussion

### The DFG motif of phosphorylated AurA is predominantly in the DFG-Out state

We set out to explain how phosphorylation of AurA on T288 leads to a ~100-fold increase in catalytic activity (Supplementary Figure S1)^19,20^. We previously used an infrared (IR) probe that tracks the DFG motif of AurA to show that Tpx2 binding triggers a conformational change from the DFG-Out to the DFG-In state^21^. In this method, a cysteine residue is introduced at position Q185 at the back of the active site of AurA, and chemical labeling is used to introduce a nitrile infrared probe at this position^25^. To test whether phosphorylation of AurA also causes a conformational shift of the DFG motif, we prepared samples of AurA Q185C phosphorylated on T288. Homogeneous phosphorylation and nitrile labeling were verified by western blotting and mass spectrometry (Supplementary Figure S2 and S3).

IR spectra of labeled phosphorylated AurA bound to ADP showed predominantly a single absorbance band centered at 2158 cm^−1^ (Figure 1a). We previously assigned this peak in spectra of unphosphorylated AurA to the DFG-Out form of the kinase, in which the nitrile probe is buried in a hydrophobic pocket (Figure 1b)^21^. Addition of saturating amounts of Tpx2 peptide (residues 1-43 of human Tpx2) to the IR samples caused a dramatic change in the spectra in which the central peak at 2158 cm^−1^ largely disappears, and two new peaks appear at 2149 cm^−1^ and 2164 cm^−1^ (Figure 1a). These changes are indicative of a shift to the DFG-In state, in which water molecules coordinated to the DFG motif form hydrogen bonds to the probe, causing pronounced spectral shifts (Figure 1b)^21^. To confirm that the peak at 2158 cm^−1^ arises from the DFG-Out state, we mutated residue W277, which is positioned directly against the IR probe in the DFG-Out state, but is displaced away from it in the DFG-in state, to alanine (Figure 1b,c). IR spectra of the W277A mutant showed a clear spectral shift of the 2158 cm^−1^ peak (Figure 1c), consistent with this peak arising from the DFG-Out state.

**Figure 1.**
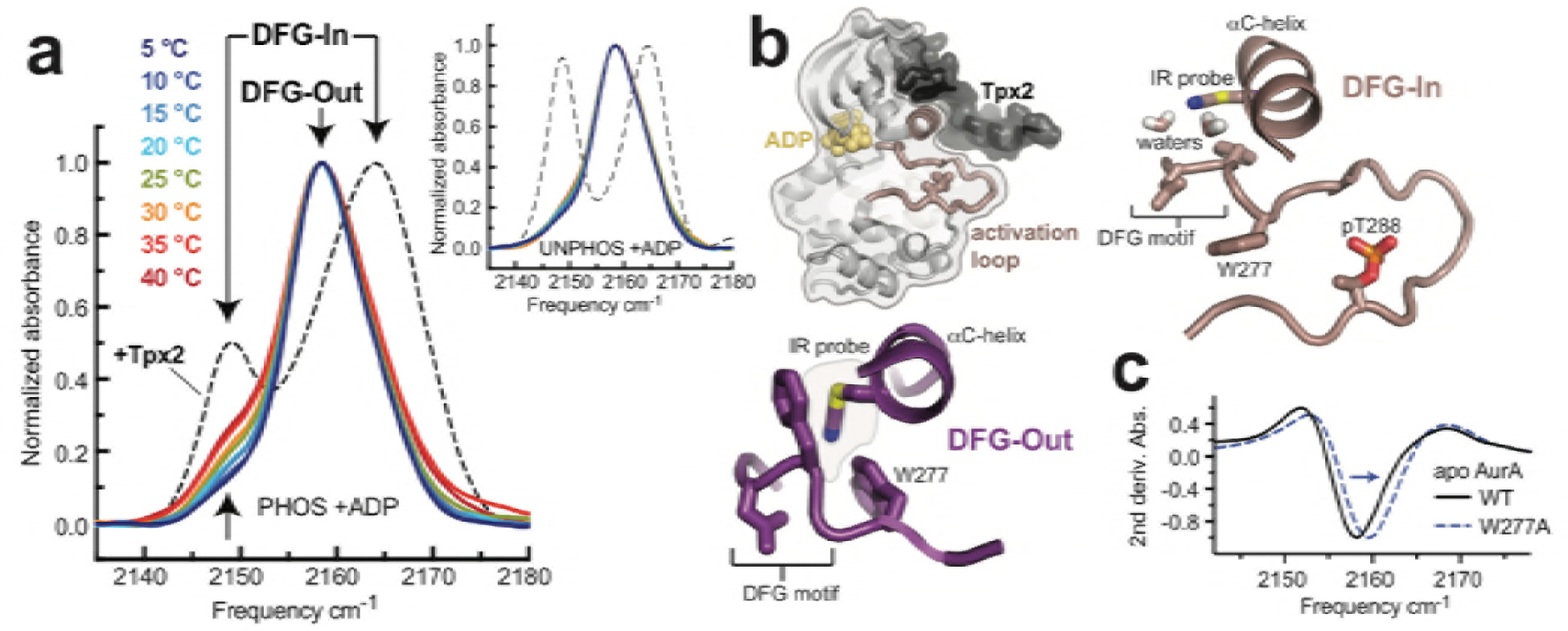
The DFG motif of phosphorylated AurA remains predominantly in the DFG-Out state. **a**) IR spectra of nitrile-labeled and phosphorylated AurA bound to ADP, measured at the indicated temperatures (colored curves). The spectrum for the same sample bound to Tpx2 is shown for comparison (dashed black line, measured at 20°C). Arrows indicate peaks assigned to the DFG-In and DFG-Out states. The inset shows the same experiments performed with unphosphorylated AurA. Single representative spectra are shown, normalized to peak maxima. (**b**) Overview of the structure of AurA in the active conformation bound to ADP (yellow) and Tpx2 (black), with enlarged views of the DFG-In (right, PDB ID: 1OL5) and DFG-Out (bottom, PDB ID: 5L8K) states with the nitrile probe (Q185CN) modeled into the structures. (**c**) Second derivatives of IR spectra of apo WT and W277A AurA, showing the spectral shift of the 2158 cm^−1^ peak (arrow).

Experiments performed over a range of temperatures highlighted the presence of a DFG-In subpopulation in the phosphorylated samples bound to nucleotide, apparent as small shoulders on either side of the main 2158 cm^−1^ peak that increase in amplitude at higher temperature (Figure 1a). A similar DFG-In subpopulation was also detected in unphosphorylated AurA bound to ADP^21^, although the temperature dependence is absent in the unphosphorylated protein (Figure 1a, inset). These surprising results show that, at physiological temperatures, phosphorylation does not significantly change the DFG-In/Out equilibrium and the DFG-In subpopulation remains low. Tpx2 binding appears to be required to elicit a shift to the DFG-In state.

### Tpx2 binding shifts the phosphorylated activation loop to a more active conformation

We used intramolecular FRET to track movements of the activation loop of AurA with and without phosphorylation on T288. Donor (D) and acceptor (A) Alexa fluorophores were incorporated on the activation loop (S284C) and αD helix (L225C) as previously described^21^. These labeling positions were chosen to track the movement of the activation loop across the active site as the kinase switches from the DFG-Out to the DFG-In state, with the dyes predicted to be further apart in the DFG-In state (Figure 2a). Phosphorylation of the protein on T288 was confirmed by tryptic mass spectrometry (Supplementary Figure S4). The labeled phosphorylated kinase exhibits robust catalytic activity in the absence of Tpx2, and is further activated only modestly by the addition of Tpx2 (3-4 fold), a characteristic feature of WT AurA phosphorylated on T288 (Supplementary Figure S5)^19,20^.

Steady-state fluorescence emission spectra were measured for D- and D+A-labeled samples in ligand titration experiments. The addition of either nucleotide or Tpx2 resulted in enhanced fluorescence emission from the donor dye and reduced emission from the acceptor, consistent with a decrease in FRET efficiency (Figure 2a). This indicates that upon binding nucleotide and Tpx2, phosphorylated AurA undergoes a conformational change to a more active conformation in which the activation loop is extended and the dyes are farther apart. The scale of the conformational change was estimated by calculating ensemble-averaged inter-dye distances from the bulk FRET efficiencies (see Methods). The maximal increase in distance is observed when the kinase is saturated with both nucleotide and Tpx2, and is on the order of ~1 nanometer for both phosphorylated and unphosphorylated AurA (Figure 2b, Supplementary Figure S6)^21^. This suggests that the activation loop undergoes a similar structural change in response to ligand binding regardless of its phosphorylation state. However, we noted several differences between the unphosphorylated and phosphorylated enzymes in how they respond to Tpx2 binding. Firstly, the affinity of the phosphorylated kinase for Tpx2, determined from the titration data, is ~20-fold higher than for the unphosphorylated kinase (Figure 2c). Secondly, phosphorylation enhances the cooperativity between nucleotide and Tpx2 binding observed in the unphosphorylated enzyme (Figure 2c, black arrows). Thirdly, Tpx2 binding alone is sufficient to produce a maximal conformational shift in the phosphorylated kinase, whereas, even at saturating concentrations, Tpx2 is insufficient to achieve this for the unphosphorylated kinase, and ADP and Tpx2 must both be present (double-headed arrow in Figure 2b, Supplementary Figure S6). These results suggest that Tpx2 and phosphorylation have synergistic effects on AurA and work together to fully stabilize the kinase in the active DFG-In state.

In contrast to the differing responses to Tpx2, AurA responds similarly to nucleotide binding regardless of the phosphorylation state of the enzyme. Specifically, the magnitude of the conformational change induced by nucleotide binding, as inferred from the increase in inter-dye distance, is similar in both cases and approximately half the maximal change (arrows in Figure 2b), and the nucleotide affinity is not increased by phosphorylation in the absence of Tpx2 (Figure 2c, right panel). Similar results were obtained with the non-hydrolyzable ATP analog AMPPNP (Supplementary Figure S6). These data suggest that the binding of nucleotide to the active site of phosphorylated AurA is only weakly coupled to the conformation of the activation loop, but that they become tightly coupled when Tpx2 is present.

**Figure 2.**
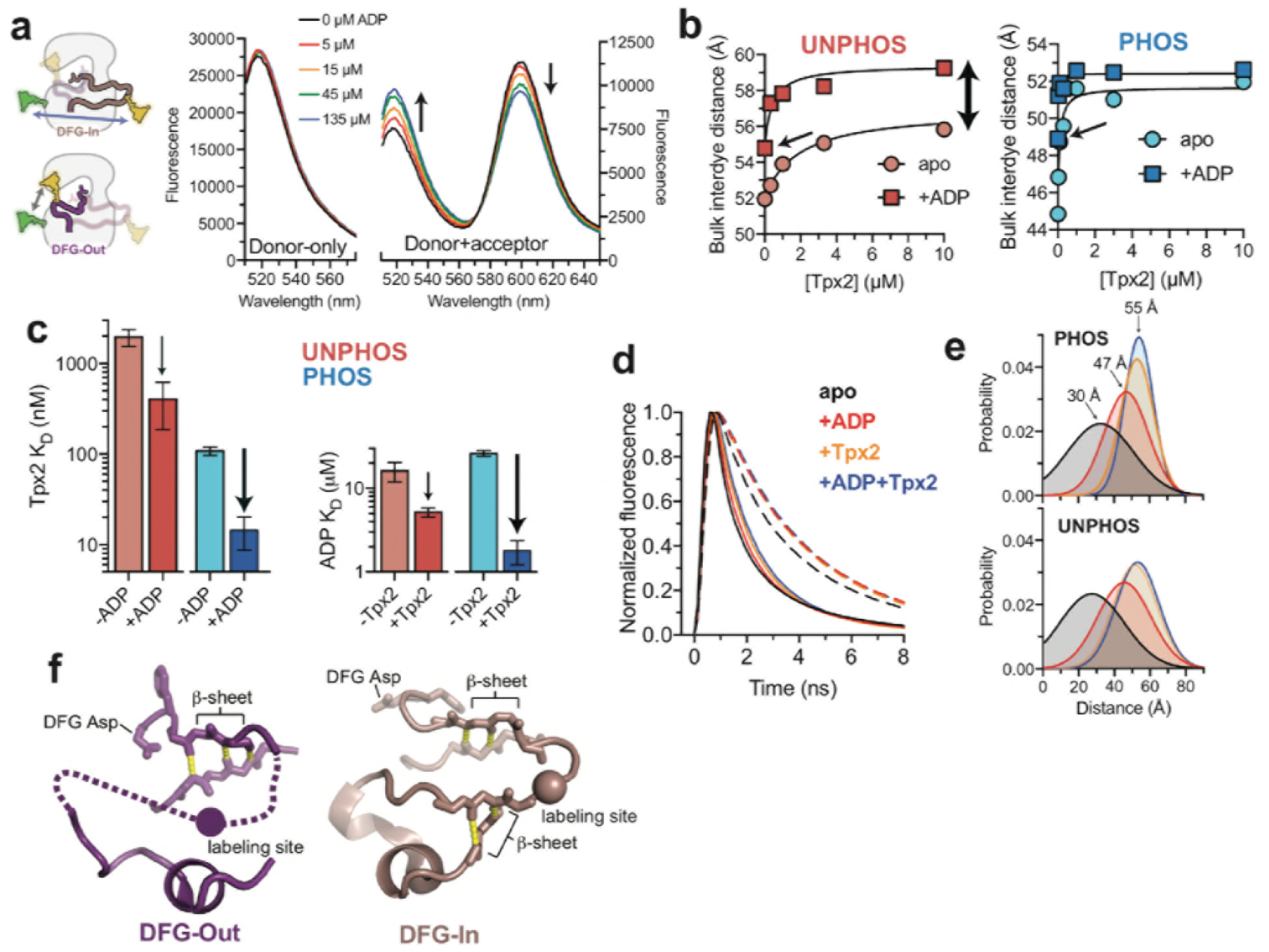
The phosphorylated activation loop remains flexible and shifts to a more active conformation upon Tpx2 binding. (**a**) (left) Schematics showing the labeling scheme used and (right) emission spectra of donor-only (D, left) and donor + acceptor (D+A, right) labeled phosphorylated AurA samples in the presence of different concentrations of ADP. Single representative experiments. (**b**) Ensemble-averaged distances between donor and acceptor dyes, calculated from bulk FRET measured for unphosphorylated (left) and phosphorylated (right) AurA with varying concentrations of Tpx2 in the presence and absence of saturating ADP. Thin arrows highlight the intermediate distances observed with saturating ADP alone, and the double-headed arrow shows the incomplete shift observed with saturating Tpx2 alone for the unphosphorylated sample. Single representative experiments are shown. (**c**) Binding constants of Tpx2 (left) and ADP (right) for phosphorylated (blue) and unphosphorylated (red) AurA in the presence and absence of the other ligand. Arrows highlight cooperativity between ADP and Tpx2. Data represent mean values ± s.d.; n = 3. (**d**) Time-resolved fluorescence waveforms for D-only (dashed lines) and D+A (solid lines) phosphorylated AurA in the presence and absence of 100 μM Tpx2 and 200 μM ADP. Data are for a single representative experiment, normalized to the fluorescence peak. (**e**) Comparison of single-Gaussian distance distribution fits to fluorescence lifetime data obtained with phosphorylated (top) and unphosphorylated AurA (bottom). Same coloring as d. (**f**) Structures of the DFG-Out (left) and DFG-In (right) states of AurA, highlighting the β-sheet hydrogen bonds constraining the N- and C-terminal segments of the activation loop. The S284C labeling site is shown as a sphere.

### The phosphorylated activation loop adopts a range of conformations but becomes highly ordered upon Tpx2 binding

The steady-state FRET measurements provide ensemble-averaged measures of distance. To gain insight into the distribution of conformations present in AurA and how it is altered by ligand binding, we performed time-resolved (TR) FRET experiments to quantify energy transfer through its effect on the fluorescence lifetime of the donor dye. TR fluorescence decays were recorded for phosphorylated and unphosphorylated AurA samples in the presence and absence of saturating ADP and Tpx2 using time-correlated single-photon counting (TCSPC) (Figure 2d). These data were then fit to a structural model consisting of a Gaussian distribution of inter-fluorophore distances for each condition^26–28^ (Figures 2e). The fraction of the D+A samples lacking acceptor dye was explicitly accounted for in the TR-FRET fitting, yielding more reliable distances than the values estimated by steady-state FRET.

The distance distributions measured for the phosphorylated and unphosphorylated kinase are strikingly similar (Figure 2e). In both cases, a broad distribution centered at ~30 angstroms is observed for apo AurA, indicating that the activation loop is highly flexible under these conditions. This is consistent with adoption of the DFG-Out state, in which the C-terminal half of the activation loop lacks contacts with the rest of the kinase domain, and is typically disordered in x-ray structures^29–31^ (Figure 2f). The addition of both ADP and Tpx2 together yields the longest distances (~55 angstroms) with the narrowest distributions, indicative of a well-defined structure consistent with the DFG-In state, in which the segment of the loop containing the labeling site is anchored to the C-terminal lobe of the kinase on both sides by backbone hydrogen bonds^22^ (Figure 2f). For the +Tpx2 samples, the presence of phosphorylation resulted in additional narrowing of the distributions, suggesting that phosphorylation further restricts the movement of the loop within the DFG-In state. In the presence of ADP alone the observed distance distributions are intermediate in both distance and width between the other samples, consistent with both unphosphorylated and phosphorylated AurA remaining in an equilibrium between DFG-Out and DFG-In states (Figure 2e).

### DEER experiments confirm that phosphorylated AurA requires Tpx2 to switch to the active state

To independently confirm that the activation loop of phosphorylated AurA samples multiple conformational states, we used double electron-electron resonance (DEER) EPR spectroscopy^32^. DEER experiments probe the distance-dependent dipole-dipole interactions of unpaired electron spins, providing information about the distribution of spin-spin distances present in the sample. Two MTSL spin labels were incorporated into AurA at the same positions used for FRET experiments (L225C and S284C; labeling and phosphorylation were confirmed by mass spectrometry (Supplementary Figure S7)), and samples were flash frozen in the presence of saturating concentrations of ADP and/or Tpx2 for DEER experiments.

Background-corrected dipolar evolution data (DEER spectra) acquired for the unphosphorylated and phosphorylated samples lacking Tpx2 were very similar, with the DEER signal decaying rapidly, consistent with phosphorylation failing to trigger a shift towards the DFG-In state, and the activation loop adopting multiple conformations (Figure 3a). The addition of Tpx2 to the phosphorylated samples resulted in striking changes (Figure 3b), with pronounced oscillations apparent in the DEER spectra that persist beyond 5 microseconds, indicating a high degree of structural order in the activation loop. Consistent with this, spin-spin distance distributions determined from these data by Tikhonov regularization^33^ are similar and broad for the samples bound to nucleotide alone (Figure 3a), but display two sharp peaks at 44 and 52 angstroms for the phosphorylated samples containing Tpx2 (Figure 3b). The 52-angstrom distance is considerably longer than the distances observed in the samples lacking Tpx2, consistent with the activation loop adopting the extended conformation characteristic of the active DFG-In state (see Figure 2f). Although the shorter 44-angstrom distance could in principle arise from a fraction of the sample occupying the DFG-Out state, we consider this unlikely. Firstly, the sharp nature of the 44- and 52-angstrom peaks is indicative of the high degree of structural order expected for the DFG-In state, but not the dynamic DFG-Out state, as discussed above (see Figure 2f). Secondly, the IR and FRET results indicate that Tpx2 shifts the phosphorylated kinase mostly to the DFG-In state.

**Figure 3.**
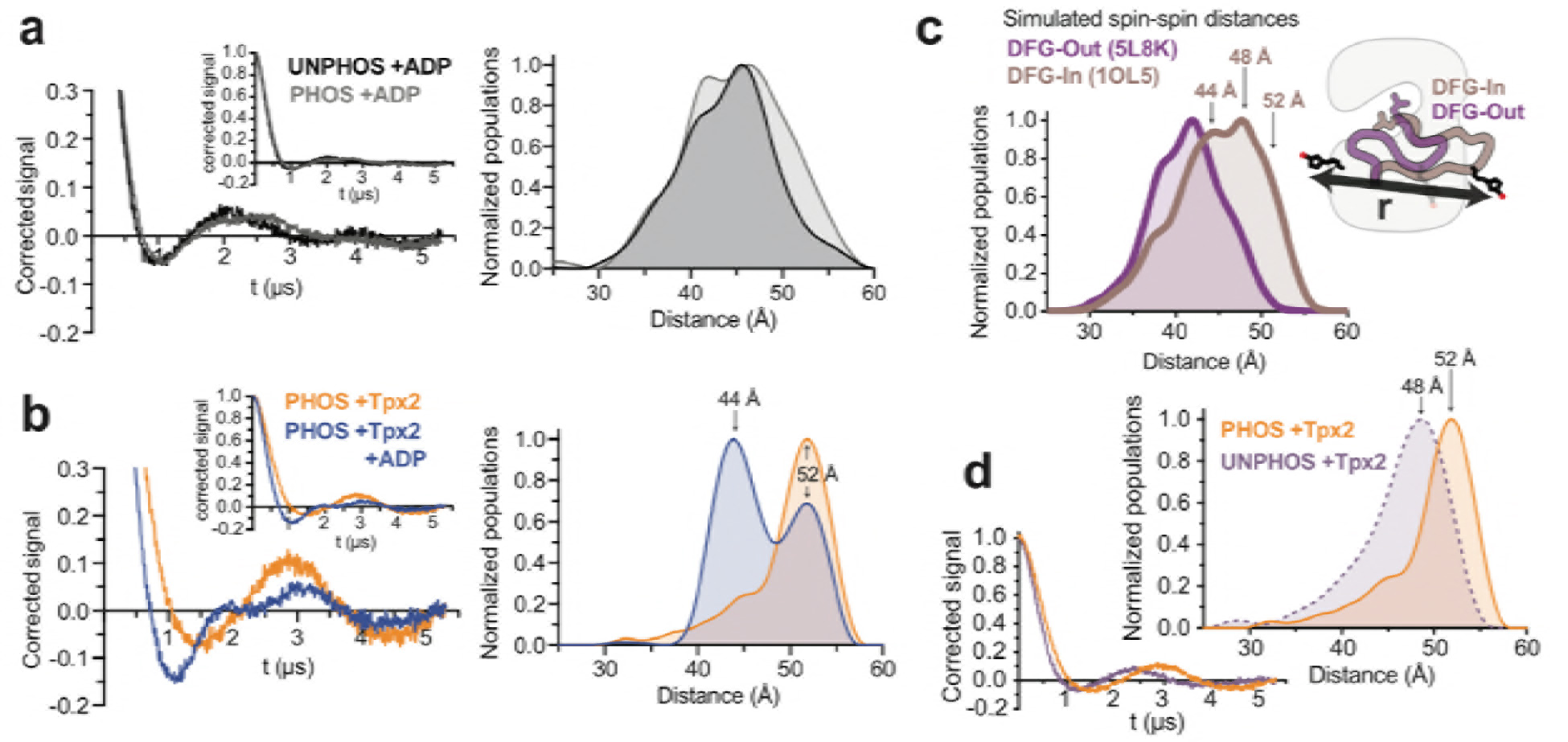
DEER spectroscopy confirms that the phosphorylated kinase requires Tpx2 to switch fully to the active state. (**a-b**) Enlarged view of the background-corrected DEER spectra are shown on the left with the full spectra shown as insets. The corresponding population densities obtained by Tikhonov regularization are shown on the right. All figures show data from a single representative experiment. (**a**) Comparison of unphosphorylated (black) and phosphorylated (gray) AurA in the presence of ADP. (**b**) Comparison of phosphorylated AurA bound to Tpx2 alone (orange), and both Tpx2 and ADP (blue). The 44- and 52-angstrom peaks in the distance distribution are highlighted. (**c**) Spin-spin distance distributions obtained by molecular dynamics simulations initiated from x-ray structures of AurA in either the DFG-Out inactive state (purple) or the DFG-In state with both Tpx2 and phosphorylation, representing the fully active conformation (pink). The inset shows a schematic of the labeled kinase. (**d**) DEER spectra and distance distributions comparing unphosphorylated (purple) and phosphorylated AurA (orange) bound to Tpx2.

To support the DEER experiments we performed molecular dynamics simulations of MTSL-labeled AurA in either the DFG-Out state (PDB ID: 5L8K) or the DFG-In state (PDB ID: 1OL5), totaling 75-110 microseconds of aggregate simulation data for each state. As expected, simulated spin-spin distances were considerably longer for the DFG-In state than the DFG-Out state (Figure 3c). In simulations of the most active state (the phosphorylated kinase in the DFG-In state bound to Tpx2), different spin label rotamers give rise to a range of distances, with peaks in the distribution apparent around ~44, ~48 and ~52 angstroms (Figure 3c). The 44- and 52-angstrom spin-spin distances are consistent with the DEER data, although it is not clear why the relative contribution of the 52-angstrom distance decreases upon addition of ADP (Figure 3b). Presumably the rotamer states corresponding to the 48-angstrom distance seen in the simulations are sparsely populated at the low temperature of the DEER experiment.

DEER spectra were also measured for AurA bound to Tpx2 but lacking phosphorylation (Figure 3d), and the corresponding distance distributions showed a sharp peak at a longer distance than in the corresponding sample without Tpx2 (see Figure 3a), consistent with a switch to the DFG-In state. Interestingly, this distance was ~4 angstroms shorter than the 52- angstrom peak measured for the phosphorylated sample in the presence of Tpx2 (Figure 3d), suggesting that phosphorylation alters the structure of the DFG-In state.

Taken together, the IR, FRET, and EPR data conclusively show that phosphorylation on T288 alone is not sufficient to shift AurA into the DFG-In state. Instead, the phosphorylated activation loop samples a range of different conformations spanning the DFG-In and DFG-Out states, and phosphorylation must drive catalytic activation of AurA by other mechanisms, perhaps by altering the structure and/or dynamics of the DFG-In subpopulation to populate catalytically competent geometries.

### Phosphorylation promotes a single functional conformation in the DFG-In state

To provide insight into how phosphorylation alters dynamics in the DFG-In state, we performed molecular dynamics simulations of the wild-type kinase. Simulations were initiated from the x-ray structure of active DFG-In AurA bound to ADP and Tpx2 (PDB ID: 1OL5)^22^, and were run in the presence and absence of Tpx2, with and without phosphorylation on T288. For each of these four biochemical states, 250 trajectories up to 500 nanoseconds in length were obtained on the distributed computing platform Folding@home, for a total of over 100 microseconds of aggregate simulation time for each biochemical state. Analysis of the DFG conformation revealed that the simulations remained predominantly in the DFG-In state (Figure S13), suggesting that the simulation time was insufficient to capture the slow conformational change to the DFG-Out state. The simulations can thus be regarded as probing the conformational dynamics of the DFG-In kinase.

The T288 phosphorylation site lies in the C-terminal segment of the activation loop, which forms an integral part of the binding site for peptide substrates (Figure 4a). In the crystal structure used to initiate the simulations, this segment of the loop appears to be stabilized by interactions between the pT288-phosphate moiety and three arginine residues: R180 from the αC helix, R286 from the activation loop, and the highly conserved R255 from the catalytic loop “HRD motif” (Figure 4a)^22^. To probe the integrity of these interactions in the simulations, and to investigate loop dynamics in their absence, we examined the distribution of distances between the Cζ atoms of either R180 or R255 and the Cβ or Cα atoms of T288 following equilibration within the DFG-In state (Figure 4b). For the simulations performed in the presence of phosphorylation, both the T288-R255 and T288-R180 distances are tightly clustered around 5-6 angstroms, confirming that the phosphate moiety is coordinated by both arginine residues throughout the majority of the trajectories (Figure 4b). In contrast, the distributions are considerably broader in the absence of phosphorylation, confirming that this segment of the activation loop remains dynamic.

**Figure 4.**
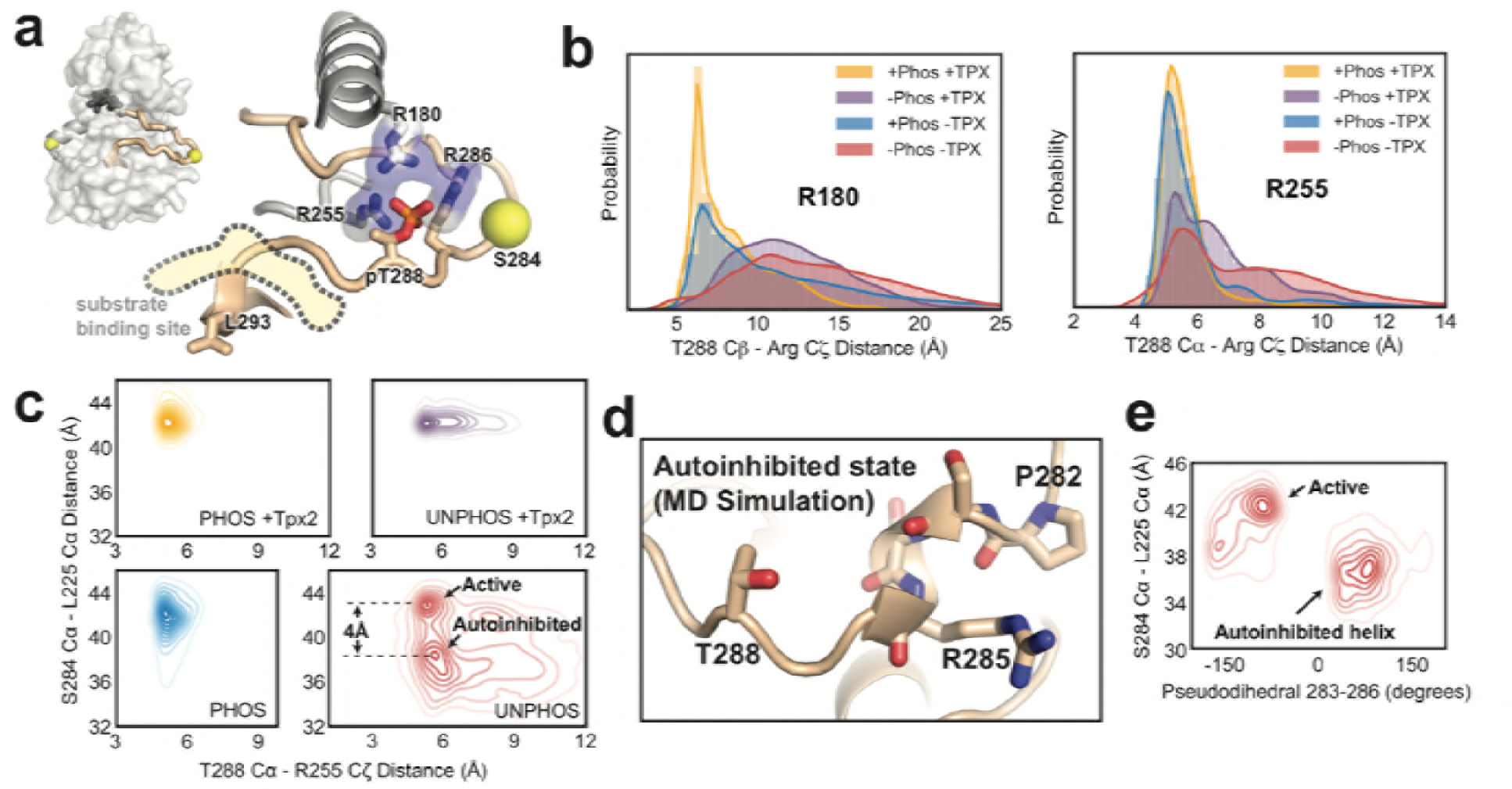
Molecular dynamics simulations of AurA in the DFG-In state show phosphorylation promotes a specific configuration of the activation loop. (**a**) Structure of active phosphorylated AurA bound to Tpx2 and ADP (PDB ID: 1OL5) showing the interactions between pT288 and the basic arginine pocket. The S284 and L225 Cα atoms are shown as spheres. (b) Distributions of the T288 Cα - R180 Cζ (left) and T288 Cα - R255 Cζ (right) distances determined from MD simulations of AurA performed in the indicated biochemical states. Note that ADP was present in all simulations. (**c**) Contour plots showing the L225 Cα - S284 Cα distances plotted against the T288 Cα - R255 Cζ distances for all four biochemical conditions. The active and autoinhibited states observed for the unphosphorylated kinase in the absence of Tpx2 (red), and the shift in the L225-S284 distance between them, are indicated. (**d**) Simulation snapshot showing the helical turn in the activation loop and the position of the T288 sidechain at the C-terminal end of the helix. (**e**) The L225 - S284 distance is plotted against the dihedral angle defined by the Cα atoms of residues 283-286 (pseudodihedral). The helical conformation in the autoinhibited state is indicated.

We also tracked the distance between the L225 and S284 Cα atoms (the sites used for incorporating spectroscopic probes) to capture movements of the tip of the activation loop containing S284 away from the active conformation. Plotting the L225-S284 distance versus the R255-T288 distance provides additional insight into the relative effects of Tpx2 and phosphorylation (Figure 4c). Phosphorylated AurA is locked into a single conformation with a long L225-S284 distance (42 Å) and short R255-T288 distance, indicative of a stable active state. Interestingly, phosphorylation alone is nearly as effective at constraining the loop in this state as phosphorylation and Tpx2 together (Figure 4c, left panels). In contrast, the simulations of unphosphorylated AurA bound to Tpx2 show a broader distribution of distances, indicating that Tpx2 is less effective than phosphorylation at stabilizing the activation loop. This may explain why Tpx2 activates AurA to a lesser extent than phosphorylation.

### Phosphorylation may disrupt an autoinhibitory DFG-In state

The simulations of unphosphorylated AurA without Tpx2 show a much greater degree of conformational heterogeneity than the simulations of the other three biochemical states. The N-terminal lobe of the kinase is particularly heterogeneous, and local unfolding occurs within the αC-helix in many of the trajectories, as seen previously in simulations of the epidermal growth factor receptor^34^ and in x-ray structures of the related AGC-family kinase Akt^35^. Although the activation loop moves substantially away from the active conformation, giving rise to shorter L225-S284 distances, the loop is not in fact disordered. Instead, two discrete subpopulations are visualized in the simulations: one subpopulation corresponding to the active-like state, and another with a much shorter L225-S284 distance (38 Å, see Figure 4c), representing a stable DFG-In state in which the activation loop is not in a catalytically-competent conformation. Manual inspection of the trajectories revealed that in this subpopulation the tip of the activation loop folds into a short helical turn spanning residues P282-R286, with the P282 proline residue serving as the N-terminal capping residue in most of the trajectories^36^ (Figure 4d). Calculating the pseudodihedral angle for the Cα atoms of S283-R286 across all trajectories confirmed that the inactive subpopulation possesses well-defined helical pseudodihedral values of 50-75° (Figure 4e). Although this conformation has not been observed in x-ray structures of AurA, the formation of short helices in the activation loop is a common feature of the inactive states of other protein kinases^37–40^.

An interesting feature of the autoinhibited DFG-In state observed in the simulations is that the T288 residue, which immediately follows the helical segment in the protein sequence, is positioned close to the C-terminal end of the helix in almost all of the trajectories (Figure 4d), with the sidechain hydroxyl forming hydrogen bonds to the backbone carbonyls of residues R285 and R286 in many of the simulation snapshots. We reasoned that upon phosphorylation of T288, the proximity of the phosphate group to the negatively-charged end of the helix dipole^41^ would destabilize this autoinhibited state, promoting the refolding of the activation loop to the active conformation. The existence of such an autoinhibition mechanism may explain why unphosphorylated AurA exhibits very low catalytic activity in the absence of Tpx2 despite a substantial DFG-In subpopulation^21^.

We wondered why the helical activation loop conformation has not been observed in x-ray structures of AurA. In fact, the activation loop adopts the active conformation in only a small subset of AurA structures determined in the presence of Tpx2^22–24^ or other protein factors that stabilize the active state^42^. Instead, almost all of the structures of AurA in the DFG-In state (76 structures out of 138 total structures of AurA in the PDB) were determined in the same hexagonal crystal form in which the kinase adopts an inactive conformation with the activation loop misaligned and the peptide binding site disassembled. Upon examination of the crystal lattice we noticed that this conformational state of the activation loop appears to be induced by a crystal contact between the peptide binding site and a neighboring molecule in the lattice (Supplementary Figure S11). This apparent crystallographic artifact may have prevented previous observation of the helical autoinhibited DFG-In state visualized in our simulations, which model the kinase in solution rather than in the crystallographic context. In conclusion, the MD simulations, which represent over a millisecond of simulation data, suggest that phosphorylation has profound effects on the activation loop of AurA in the DFG-In state, disrupting an autoinhibited state and promoting an alternative conformation primed for catalytic function.

### Experimental evidence for ordering of the activation loop in the DFG-In state by phosphorylation

A key insight from the MD simulations is that phosphorylation alone is sufficient to promote the active configuration of the activation loop, and that coordination of the phosphothreonine by the R180 and R255 residues is likely important for this. To test the role of these interactions, we mutated the R180 residue to an alanine in the context of our phosphorylated FRET construct (phosphorylation was confirmed by mass spectrometry, see Supplementary Figure S8). The R180A mutant possessed 4-fold lower activity in the absence of Tpx2, whereas the activity in the presence of Tpx2 was only modestly affected (Figure 5a), indicating that the pT288-arginine interactions are particularly important for activation of AurA by phosphorylation alone. Steady-state FRET experiments on the R180A mutant showed broadly similar ligand-induced conformational shifts as observed in the absence of the mutation, but the response to Tpx2 alone was somewhat smaller and more similar to unphosphorylated AurA^21^ (Figure 5b and Supplementary Figure S6). Consistent with this observation, the binding affinities for Tpx2 and ADP are reduced by the R180A mutation, and are similar to those observed for unphosphorylated AurA (Figure 5c). This indicates that the pT288-arginine interactions are necessary for the synergy between phosphorylation and Tpx2 observed with the wildtype enzyme.

**Figure 5.**
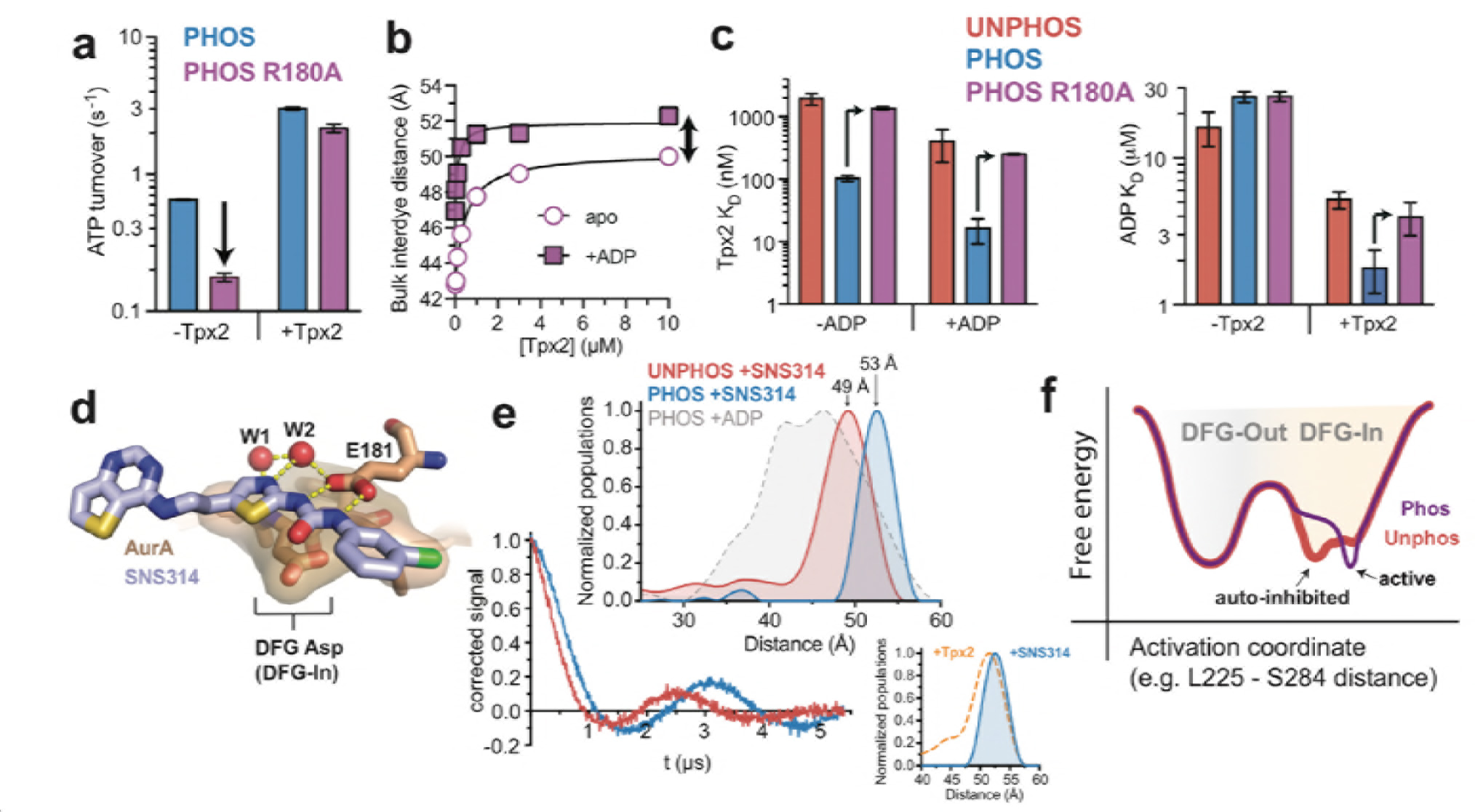
Experimental support for ordering of the activation loop in the DFG-In state mediated by phosphorylation. (**a**) Kinase activity (shown as ATP turnover per second) for phosphorylated WT (blue) and phosphorylated R180A (purple) AurA unlabeled FRET constructs in the presence and absence of 10 μM Tpx2. The decrease in the activity in the absence of Tpx2 due to the R180A mutation is highlighted by the arrow. Data represent mean values ± s.d.; n = 3. (**b**) Ensemble-averaged distances between donor and acceptor dyes, calculated from bulk FRET, for phosphorylated R180A AurA with varying concentrations of Tpx2 in the presence and absence of saturating ADP. The double-headed arrow highlights the incomplete shift observed with saturating Tpx2 in the absence of ADP. Single representative experiments are shown. (**c**) Binding constants for Tpx2 (left) and ADP (right) for phosphorylated (blue), phosphorylated R180A (purple) and unphosphorylated (red) unlabeled AurA FRET constructs in the presence and absence of saturating concentrations of the other ligand. Arrows highlight the effects of the R180A mutation. Data represent mean values ± s.d.; n = 3. (**d**) X-ray structure of SNS-314 bound to AurA highlighting interactions with the DFG motif, structured water molecules and the catalytic glutamate (E181) that promote the DFG-In state. (**e**) DEER spectra (bottom) and distance distributions (top) measured for unphosphorylated and phosphorylated AurA bound to SNS-314. The distribution measured for phosphorylated AurA bound to ADP is shown in the top panel for comparison. The inset shows a comparison of the distributions obtained for the phosphorylated kinase bound to either SNS-314 (blue) or Tpx2 (orange), highlighting their similarity. (**f**) Hypothesized energy landscape for AurA, highlighting the effect of phosphorylation on the DFG-In state.

The catalytic defect of the R180A mutant suggests that the pT288-phosphate moiety does dock into the arginine pocket even when Tpx2 is absent, as observed in the MD simulations. We hypothesized that our FRET and EPR experiments did not detect this in the form of a conformational change in the activation loop (see for instance Figure 3a) because under these conditions the substantial DFG-Out subpopulation masks the structural changes occurring in the DFG-In subpopulation. To test this, we used the ATP-competitive AurA inhibitor SNS-314, which preferentially binds to the DFG-In state of AurA^43^ (Figure 5d), to induce a homogeneous population of DFG-In kinase. DEER spectra measured on unphosphorylated and phosphorylated AurA bound to SNS-314 were strikingly different from one another, and the Tikhonov distributions confirmed that phosphorylation causes a pronounced shift of ~4 angstroms to longer distance (Figure 5e). Both distance distributions are very narrow, consistent with the activation loop adopting a well-defined structure in the absence of phosphorylation that differs from that of the active state. The increase in spin-spin distance upon phosphorylation is similar in magnitude to the change in the L225-S284 Ca distance between the autoinhibited and active DFG-In states in the MD simulations, and may correspond to this conformational change (Figure 4c). Importantly, the distance measured for phosphorylated AurA bound to SNS-314 is nearly identical to that observed for the phosphorylated kinase bound to Tpx2 (Figure 5e inset), the most catalytically active form of the enzyme that x-ray structures show to be in the fully active state.

These results confirm that phosphorylation is both necessary and sufficient to fully constrain the activation loop in the active conformation for the fraction of the kinase adopting the DFG-In state. We conclude that while phosphorylated AurA samples both the DFG-Out and DFG-In states, the structure and dynamics of the DFG-In subpopulation are profoundly altered by phosphorylation, leading to catalytic activation of the kinase. The autoinhibited DFG-In state identified in the MD simulations provides an explanation for how phosphorylation can promote activity without triggering a shift in the DFG equilibrium (Figure 5f). In this model, phosphorylation acts as much by destabilizing the autoinhibited DFG-In state as by stabilizing the active DFG-In state, leading to a population shift within the DFG-In state without much effect on the relative populations of the DFG-In and DFG-Out states.

### A speculative model for a role of the DFG flip in nucleotide release during catalytic turnover of AurA

We wondered whether maintaining a significant DFG-Out population might be important for the physiological function of AurA. In the closely-related kinase PKA catalytic turnover is rate-limited by product dissociation^44–46^, and it has been previously suggested for other kinases that the DFG flip may be coupled to nucleotide binding and release^47,48^. We therefore performed rapid mixing experiments to measure the rate of ADP binding to fluorescently labeled AurA, using a transient time-resolved fluorescence instrument that can track structural changes by time-resolved FRET with millisecond time resolution^49^.

For both unphosphorylated and phosphorylated samples lacking Tpx2 a single kinetic phase was observed in the mixing experiments (Figure 6a). Fitting of the time-resolved waveforms for the phosphorylated sample yielded FRET distances that evolved from ~37 angstroms at early time points to ~47 angstroms at the end of the mixing experiment, demonstrating that the experiment monitors the shift towards the DFG-In state triggered by nucleotide binding (Figure 6a,b). A linear dependence of the apparent association rate constant, k_obs_, on ADP concentration was measured for both samples (Figure 6c), consistent with the kinetics reporting directly on nucleotide binding. In the presence of Tpx2 the kinetic behavior changed dramatically, with both the unphosphorylated and phosphorylated samples exhibiting two well-separated kinetic phases (Figure 6d, Figure S9), which were analyzed separately as discussed in more detail in the Supporting Information. The observed rate constants for the fast phase, k_FAST,obs_, were linearly dependent on ADP concentration (Figure 6e), consistent with ADP binding, whereas the slow phase was not strongly dependent on ADP concentration (Figure S10), suggesting a structural change.

**Figure 6.**
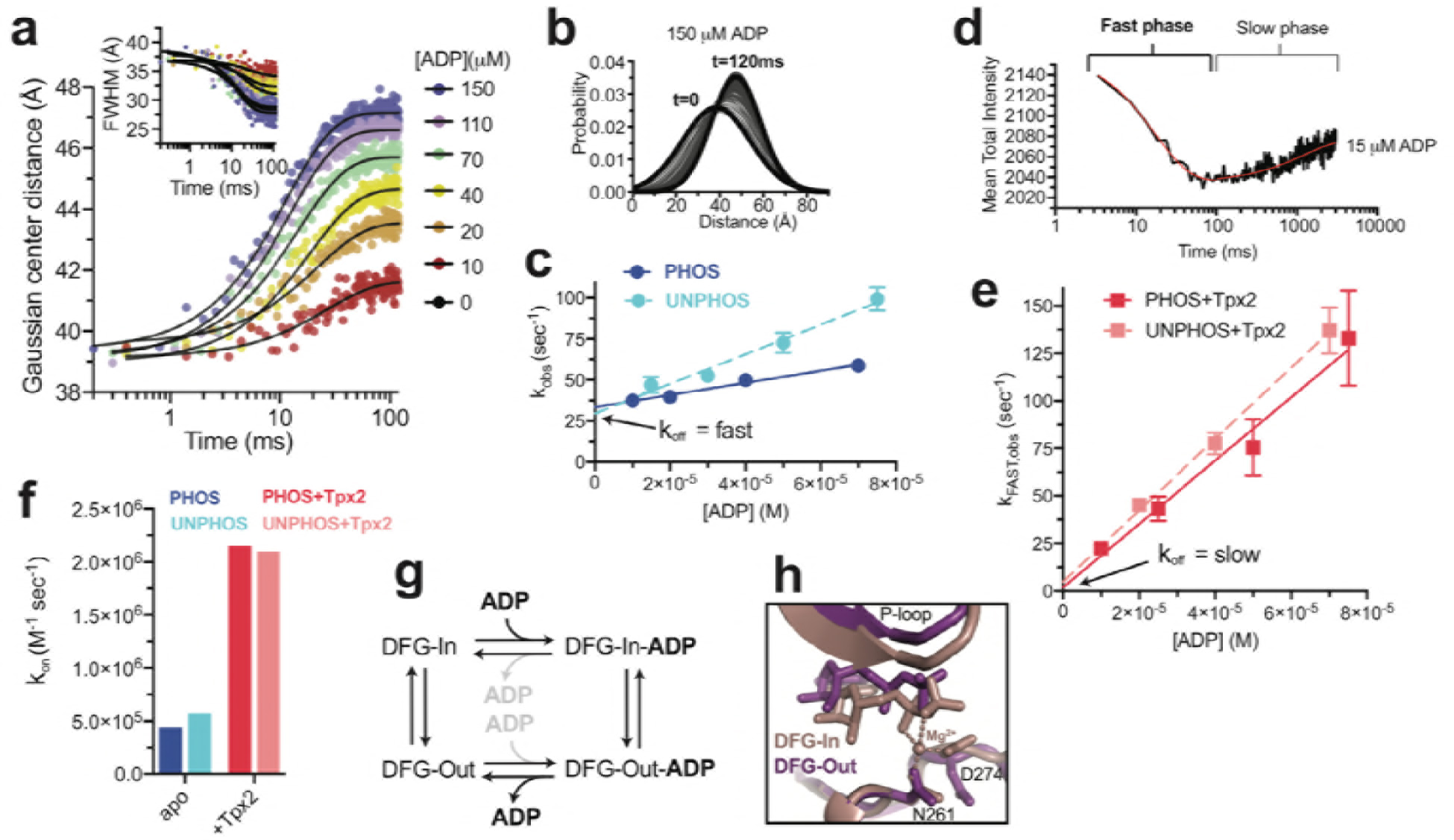
Tpx2 enhances nucleotide binding kinetics and slows nucleotide release. (**a**) Gaussian FRET distances (main graph) and corresponding full-width half maximum values (inset) determined by transient time-resolved FRET for phosphorylated AurA mixed with indicated concentrations of ADP. Black lines are single exponential fits. Each trace represents a single experiment. (**b**) The distance distributions derived from the lifetime data are shown for a single injection experiment for time points between 0 and 120 milliseconds. (**c**) Apparent rate constants k_obs_ determined from single exponential fits to the fluorescence data, plotted as a function of [ADP] for phosphorylated (dark blue) and unphosphorylated (light blue) samples. Data represent single [ADP] series experiments, and error bars represent the estimated errors from linear regression. The arrow highlights the measured off rates of ~30 s^−1^ for both samples. (**d**) Fluorescence transient for unphosphorylated AurA bound to Tpx2 and mixed with 15 μM ADP. The fast and slow phases are indicated. (**e**) Rate constants k_fast,obs_ determined from the fast phase are plotted as a function of [ADP]. (**f**) Elementary rate constants k_ON_ for ADP association in the presence and absence of 20 μM Tpx2. Data represent the average of two independent experiments. (**g**) Kinetic scheme for ADP binding to AurA, emphasizing the proposed model in which ADP binds preferentially to the DFG-In state and dissociates from the DFG-Out state. (**h**) Comparison of x-ray structures of AurA bound to ADP in the DFG-In (pink, PDB ID: 1OL5) and DFG-Out (purple, PDB ID: 5L8K) states, with the magnesium coordination indicated.

We propose the following interpretation of these results (see Supporting Information for further discussion). The DFG-In and DFG-Out states are in equilibrium, and nucleotide binds preferentially to the DFG-In subpopulation. In the absence of Tpx2, the DFG flip is fast compared to binding resulting in a single kinetic phase in which the relatively slow on rate k_ON_ (~4 ×10^5^ M^−1^s^−1^) reflects binding to the small DFG-In subpopulation. In the presence of Tpx2, the same kinetic scheme applies, but the DFG flip slows dramatically to ~2 s^−1^ (the slow phase in Figure 6d) so that the DFG-In and DFG-Out states now interconvert slowly compared to nucleotide binding. The on and off rates determined from the fast phase (Figure 6e) therefore reflect nucleotide binding and dissociation from the kinetically isolated DFG-In subpopulation. The ~5-fold increase in the on rate k_ON_ observed with Tpx2 (~2 ×10^6^ M^−1^s^−1^, Figure 6f) is consistent with the much larger DFG-In subpopulation under those conditions. The off rate k_OFF_ (the intercept in Figure 6e) is difficult to measure precisely due to the steep concentration dependence, but is clearly much slower than in the absence of Tpx2 (compare arrows in Figure 6c and e). This indicates that nucleotide dissociation from the DFG-In state is very slow, and suggests that the DFG-Out state is responsible for the fast dissociation observed in the absence of Tpx2 (30 s^−1^), and may even be necessary for efficient nucleotide dissociation (Figure 6g). This hypothesis is consistent with x-ray structures of DFG-Out AurA bound to nucleotides^19,50^, which show that magnesium coordination is lost and the nucleotide is partly dissociated from the C-terminal lobe of the kinase (Figure 6h).

In our model, the slow phase observed in the presence of Tpx2 arises from the DFG-Out subpopulation binding nucleotide and converting to the DFG-In state. FRET analysis of the lifetime data for the slow phase suggested a small shift of ~1-2 angstroms to longer distance, consistent with a small subpopulation undergoing the DFG flip (see Supporting Information and Figure S10), but the limited size of this change prevents us from ruling out other possible explanations, and our model therefore remains speculative. Interestingly, the observed timescale of this structural change (~2 s^−1^) is similar to the maximum catalytic turnover rate observed for Tpx2-bound AurA (~3 sec^−1^, Figure 5a), raising the possibility that the DFG flip is a rate limiting step for catalytic turnover under these conditions. We propose that the DFG-Out state represents an intermediate in the catalytic cycle, and that an important function of the DFG flip in AurA may be to promote efficient nucleotide exchange.

## Discussion

The majority of eukaryotic protein kinases are activated by phosphorylation on the activation loop^3^. X-ray structures have suggested that the functional role of phosphorylation is to trap the kinase in the active DFG-In state and rigidify the flexible activation loop in a specific configuration that promotes catalysis and substrate binding^4–6^. Our results show that phosphorylation can drive catalytic activation of a protein kinase without restraining the activation loop in the DFG-In state, providing a contrasting and highly dynamic view of an activated kinase in which major conformational changes of catalytic elements may occur continuously during the catalytic cycle. A recent single-molecule fluorescence study also reported that phosphorylated AurA dynamically transitions between multiple structural states^51^.

We previously reported that the binding of Tpx2 to unphosphorylated AurA causes a population shift towards the DFG-In state, in striking contrast with the phosphorylation-mediated activation mechanism described here^21^. Our simulation and EPR data also reveal differences in how phosphorylation and Tpx2 affect the DFG-In subpopulation, with Tpx2 failing to fully constrain the C-terminal segment of the activation loop in the active conformation, unlike phosphorylation. Thus it appears that, when acting alone, phosphorylation and Tpx2 activate AurA through distinct pathways, with phosphorylation promoting a specific configuration of the activation loop for the DFG-In subpopulation, and Tpx2 instead triggering a DFG flip as well as stabilizing the kinase αC-helix and an associated water-mediated hydrogen bond network^21^. When AurA is simultaneously activated by both Tpx2 and phosphorylation, the two regulatory inputs synergize, with the kinase switching fully to the DFG-In state and the activation loop becoming highly ordered. This synergy likely accounts for the observation that Tpx2 and phosphorylation together can overcome the deleterious effects of destabilizing mutations in the regulatory spine of AurA^21^. Interestingly, this doubly-activated form of AurA is not thought to occur in normal cells, but is prominent in about 10% of melanoma patients, where mutational inactivation of the PP6 phosphatase leads to accumulation of phosphorylated AurA on the mitotic spindle, and results in genomic instability^52,53^. The synergistic action of phosphorylation and Tpx2 on the conformational dynamics of AurA may provide unique opportunities for the development of inhibitors that selectively target this pathological form of the kinase in melanoma^54^.

Although activation of AurA by phosphorylation and Tpx2 are normally mutually exclusive, similar activation steps must occur in concert in the related AGC-family kinases, which require both phosphorylation and the docking of their C-terminal hydrophobic motifs (HM) to the αC-helix, which resembles the Tpx2 interaction. The suppressed dynamics of doubly-activated AurA may therefore be representative of canonically-activated AGC kinases, and the highly dynamic nature of phosphorylated AurA in the absence of Tpx2 may be a unique property of the Aurora kinases that reflects the loss of the HM and the emergence of the dual activation mechanisms in this lineage^21^. These dynamics may facilitate further regulation of AurA by additional cellular factors, allowing for graded levels of catalytic activity. For instance, phosphorylated AurA is known to interact with Cep-195^55^, Bora^14,15^, and Ajuba^56^ at the centrosome, and these interactions can further regulate AurA activity towards specific substrates.

The DFG flip has long been considered one of the key regulatory mechanisms used by nature to control the catalytic activity of protein kinases^7,8^. Although AurA does adopt the DFG-Out state, our results show that activation of the enzyme by phosphorylation is not mediated by a DFG flip, but rather by tuning the catalytic activity of the DFG-In subpopulation. This suggests that evolution has moderated the ionic interactions between the phosphothreonine residue and the basic residues it interacts with to ensure a substantial DFG-Out subpopulation. Based on our kinetic studies, we speculate that this may be due to the importance of the DFG-Out state for promoting nucleotide release during the catalytic cycle. If this model is correct, the surprising result reported here that phosphorylation is not tightly coupled to the DFG equilibrium may simply be the consequence of evolutionary selective pressure to optimize the kinetics of substrate binding and product release for efficient catalytic turnover. It remains to be seen whether a large DFG-Out subpopulation is a unique feature of AurA, or a more general property of activated protein kinases.

## Acknowledgements

We thank LeeAnn Higgins, Todd Markowski and Joseph Dalluge for help with mass spectrometry experiments. We thank Tanya Freedman for critical reading of the manuscript, and Wendy Gordon for helpful discussions. This work was supported by NIH grants GM102288, CA217695 (N.M.L.), F32GM120817 (E.F.R.) and P30-CA008748 and GM121505 (J.D.C.). J.D.C, J.M.B., S.M.H., and S.K.A acknowledge support from the Sloan Kettering Institute.

## Author contributions

E.F.R. and N.M.L conceived and designed experiments; E.F.R. prepared fluorescently labeled samples and performed steady-state fluorescence experiments; E.F.R. and J.M.M. performed time-resolved fluorescence experiments and analyzed the data; E.F.R. and E.L. prepared MTSL-labeled samples and A.T. performed EPR experiments and analyzed the data; D.D.T. helped conceive the EPR experiments and supervised analysis of the time-resolved FRET and EPR data; S.C. prepared nitrile-labeled IR samples and performed IR experiments; J.D.C. and N.M.L. conceived and designed molecular dynamics simulations; J.M.B. performed the MD simulations of WT AurA, S.K.A. performed the simulations of spin-labeled AurA, and S.K.A. and S.M.H. analyzed MD simulation data; E.F.R. and N.M.L. wrote the manuscript.

## Materials and Methods

### Expression and purification of AurA constructs

Aurora A kinase domain constructs were expressed and prepared as previously described^21^. Site-directed mutagenesis using the QuikChange Lightning kit (Agilent) was used to incorporate mutations R180A and W277A. Homogeneous phosphorylation of our Cys-lite constructs was achieved using a C290A variant which autophosphorylates efficiently on T288 when expressed in E.coli^57^. Phosphorylation was verified by mass spectrometry and activity assays (see Supplemental Information).

### IR spectroscopy

Phosphorylated Q185C AurA protein samples (human AurA residues 122-403 containing an N-terminal hexahistidine tag) were prepared using a cysteine-light form of the kinase in which all endogenous solvent-accessible cysteines were removed by mutagenesis (Q185C, C247A, C290A, C393S). After Nickel-affinity purification, repeated rounds of cation exchange chromatography were used to isolate the homogeneous singly phosphorylated species, with enrichment of the phosphorylated species tracked during purification by western blotting and activity assays (Supplementary Figure S2). Nitrile labeling of the purified protein was performed using a 1.5:1 molar ratio of DTNB (Ellman’s reagent), followed by 50 mM KCN, and excess labeling reagents were removed using a fast desalting column (GE Healthcare). Incorporation of a single nitrile label was confirmed by whole-protein mass spectrometry (Supplementary Figure S3). Samples for IR spectroscopy were prepared by concentrating labeled protein (50-100 μM) in the presence or absence of 4 mM ADP and 8 mM MgCl_2_, and/or excess Tpx2 peptide (~150 μM, residues 1-43 of human Tpx2, Selleckchem) in FTIR buffer (20 mM Hepes, pH 7.5, 300 mM NaCl, 20% glycerol). Samples were concentrated to ~1 mM and loaded into a calcium fluoride sample cell mounted in a temperature-controlled housing (Biotools) for IR experiments. IR spectra were recorded on a Vertex 70 FTIR spectrometer (Bruker) equipped with a liquid nitrogen cooled indium antimonide detector with 2 cm^−1^ spectral resolution. Spectra were averaged across 256 scans, background subtracted using absorbance spectra of the buffer flow-through from sample concentration, and baselined using the polynomial method in the OPUS software (Bruker).

### Kinase assays

Kinase activity was measured using the ADP Quest coupled kinase assay (DiscoverX) in a fluorescence plate reader (Tecan Infinite M1000 PRO) as described previously^21^. Assays were performed using 2, 5, 10, 100, or 200 nM kinase (depending on the protein variant), 1 mM kemptide peptide substrate (Anaspec), 10 μM Tpx2 residues 1-43 (Selleckchem), and 50 μM ATP (Sigma Aldrich). Activity was determined using the initial fluorescence intensity slopes as a function of time (determined by linear regression) for ADP concentrations 1-10 μM. Background ATPase activity was determined for samples with no peptide substrate added, and was subtracted from the activity in the presence of kemptide. We then determined the average fluorescence over the time of the assay for known ADP concentrations in the dynamic range of the assay to construct a calibration curve, and used this to convert the background-corrected activity to ATP turnover numbers. Activities given are the average of three experiments, where error bars are the standard deviation of the replicates.

### Fluorescence and Förster resonance energy transfer (FRET) experiments

For FRET experiments, the variant AurA C290S/A C393S L225C S284C was expressed, purified and labeled with donor (Alexa 488, Invitrogen) and acceptor (Alexa 568) using cysteine-maleimide chemistry, as previously described^21^. Labeled samples were validated using activity assays and mass spectrometry and retained close to full kinase activity (Supplementary Figures S3 and S4).

#### Steady-state FRET

Ligand titration FRET experiments were performed in a Fluoromax 4 Spectrofluorometer (Horiba) at 22 °C. Assays were performed in 15 mM HEPES pH 7.5, 20 mM NaCl, 1 mM EGTA, 0.02% Tween-20, 10 mM MgCl_2_, 1% DMSO and 0.1 mg/mL bovine-γ-globulins at AurA concentrations of 5-50 nM. Bulk FRET efficiency and inter-fluorophore distance were calculated from the ratios of the donor fluorescence in the presence and absence of acceptor, assuming a value of 62 angstroms for the Förster radius, as previously described^21^. The steady-state anisotropy was below 0.2 for all samples and did not change appreciably between biochemical conditions (Supplementary Figure S12).

Dissociation constants K_D_ were determined using the spectra obtained with the D+A-labeled sample. The ratio F_D_/F_A_ (where F_D_ is the donor fluorescence maximum and F_A_ is the acceptor fluorescence maximum), a highly sensitive measure of ligand binding, was fit as a function of ligand concentration. For calculation of the K_D_ for Tpx2 with saturating ADP bound, K_D_ is near the concentration of fluorescent protein. Ligand depletion was accounted for by fitting the raw data to determine the plateau F_D_/F_A_ value (representing ligand saturation), calculating the percent saturation for each total ligand concentration, and then back calculating the concentration of free ligand. F_D_/F_A_ was then re-fit as a function of free ligand concentration.

#### Time-resolved (TR) FRET

The instrument used to collect time-resolved fluorescence at equilibrium has been previously described^27^. Data were detected by time-correlated single photon counting. The instrument response function was obtained with the emission polarization set at vertical, while fluorescence data were collected with the emission polarization set at 54.7^°^, with a GFP band pass filter in place (Semrock).

Experimental buffer contained 15 mM HEPES pH 7.5, 20 mM NaCl, 1 mM EGTA, 0.02% Tween-20, 10 mM MgCl_2_. Experiments were performed at 100-200 nM unphosphorylated or phosphorylated FRET-labeled AurA, in the presence and absence of 100 μM Tpx2 and 200 μM ADP; one phosphorylated AurA experiment also contained 1 mM DTT. For both unphosphorylated and phosphorylated AurA, two independent experiments were performed and analyzed.

Data fitting was performed as previously described^26^. Briefly, time-resolved fluorescence waveforms were fit using custom software designed for analysis of time-resolved fluorescence^49^. The instrument response function and the model of the fluorescence decay were convolved to fit the measured time-resolved fluorescence waveform. Donor-only fluorescence waveforms were modeled using a multiexponential decay function, which accounts for the intrinsic lifetimes of Alexa 488, and two exponentials were required to fit the Alexa 488 fluorescence decay. Donor + acceptor (D+A) waveforms were modeled from the amplitudes and lifetimes present in the matched donor-only sample and modified so that a distance-dependent resonance energy transfer term, corresponding to a Gaussian distribution of inter-probe distances, describes the decrease in fluorescence lifetime relative to the donor-only control. The mean distance and full-width half maximum of the Gaussian functions were fit individually for each D+A and D−O pairing, while the parameters that described general conditions of the experiment common among all samples, such as the fraction of a given D+A sample containing D-only protein, were globally linked.

#### Stopped-flow fluorescence kinetics

The transient time-resolved fluorometer used has been previously described^28,49^ and uses a *Biologic USA* SFM/20 single-mix stopped-flow instrument coupled to a transient time-resolved fluorescence spectrophotometer based on direct waveform recording technology. The flow rate was 8 mL/sec, the instrument dead time is >2 ms, and 3-5 waveforms were averaged every 1 ms. At each ADP concentration, 12-20 successive replicate mixing experiments were performed and averaged together, and these averaged waveforms were fit to extract time-resolved FRET distance information and kinetic constants.

For experiments, 20-50 nM phosphorylated or unphosphorylated FRET-labeled AurA was loaded into syringe A, 10-300 μM ADP was loaded into syringe B, and samples were rapidly mixed. For measurements of ADP binding in the presence of Tpx2, 20 μM Tpx2 was added to the buffer in both syringes A and B. Stopped-flow experiments were performed at 25 ^°^C in buffer containing 20 mM HEPES pH 7.4, 200 mM NaCl, 10% glycerol, 10 mM MgCl_2_, 0.1 mg/mL bovine-γ-globulins, and 1 mM DTT. The kinetic constants plotted are the average obtained in two experiments.

For details of the kinetic fitting, see Supplementary Information.

### EPR experiments

DEER samples were prepared in the Cys-lite mutant construct AurA C290S/A C393S C247A L225C S284C, purified as described above. AurA was labeled with MTSL (Santa Cruz Biotechnology), purified by cation exchange chromatography, and concentrated. Labeling was verified using mass spectrometry, and MTSL-labeled samples retained close to full activity of unlabeled AurA in the presence and absence of Tpx2 (Supplementary Figure S7). The protein was then buffer exchanged into the experimental buffer, which was 20 mM HEPES pH 7.5, 300 mM NaCl, 10% deuterated glycerol, 2% v/v H_2_O in D_2_O. For DEER experiments, samples containing 30-60 μM MTSL-labeled AurA were prepared in the presence and absence of 100-200 μM Tpx2 and 300 μM ADP (8 mM MgCl_2_ was added to samples containing ADP). Final samples varied in v/v H_2_O concentration from 2-14%; however, no significant differences were observed in Tikhonov distributions derived from experiments performed in protonated and deuterated buffers. Samples were flash-frozen in an isopropanol dry ice bath followed by liquid nitrogen. Data shown are from one of two replicate experiments.

DEER spectra were detected at 65 °K using an Elexsys E580 Q-Band spectrometer (Bruker Biospin) equipped with an ER 5107D2 resonator (Bruker Biospin) using the standard 4-pulse pulse sequence with π/2 and π pulses (including ELDOR) set to 16 and 32 ns respectively. The pump frequency was set to the central resonance position of the nitroxide echo-detected field swept spectrum while the observe position was set 24G up-field to avoid excitation bandwidth overlap^27^.

Data were analyzed using custom software written in Mathematica which was based heavily on DeerAnalysis 2017^58^. The raw spectra were phase and background corrected assuming a homogeneous background model to produce the DEER waveform. Distance distributions were determined using Tikhonov regularization, with an optimal smoothing parameter chosen using a combination of the l-curve and leave one out cross validation (LOOCV) techniques^59^. After choice of smoothing parameter, a range of background fits were performed to identify stable populations in the distance distributions, with highly unstable, long distance populations being largely attributable to errors in the background fit and model choice^60^.

### Molecular dynamics simulations

#### System Preparation

*Modeling WT unphosphorylated AurA.* WT AurA in complex with ADP was simulated with and without Tpx2. All simulations were started from the x-ray structure of WT AurA bound to Tpx2 and ADP in the presence of three magnesium ions (PDB ID: 1OL5^22^). From the crystal structure, PDBFixer (https://github.com/pandegroup/pdbfixer) version 1.2 was used to model in Tpx2 residues 23-29 (unresolved in 1OL5), add hydrogens belonging to standard dominant protein residue protonation states at pH 7.4, and remove phosphorylation from threonine residues 287 and 288^61^. Crystallographic waters were retained to prevent nonphysical collapse of hydrophilic pockets during minimization. The chain containing Tpx2 was then removed for simulations without Tpx2 (−Tpx2) and retained for simulations with Tpx2 (+Tpx2). Sulfate ions present in the crystal structure were manually removed. The crystallographic ADP (containing only heavy atoms) was extracted from the structure and converted to a protonated Tripos mol2 file using OpenEye toolkit OEChem v2015.June^62,63^. The protein structure was then loaded as an OpenMM version 7.0.1 Modeller object, and the protonated ADP was reintroduced through conversion from mol2 to OpenMM format via MDTraj 1.4.2^61,64^.

*Modeling WT phosphorylated AurA.* Simulations of phosphorylated WT AurA in complex with ADP were prepared as above, but the phosphothreonines (denoted TPO in the PDB file) at positions 287 and 288 were left in place and parameterized using the Sticht T1P AMBER parameters^65^ retrieved from the AMBER parameter database^66^. The AMBER phosphothreonine parameter file was converted to OpenMM ffxml using a python script that has been made publically available (https://github.com/choderalab/AurA-materials), and subsequently converted into a hydrogen specification file for OpenMM's Modeller by hand. The PDB file generated by PDBFixer was loaded into an OpenMM Modeller object where the hydrogens and bonds were added to the TPO residues using a Forcefield object instantiated with the AMBER99Bildn parameters as well as the custom TPO parameters described above. All crystallographic waters, ADP, sulfate ions and magnesium ions were handled as with the unphosphorylated WT AurA simulations.

*Parameterization the WT AMBER simulations.* An OpenMM ForceField was instantiated using AMBER99SBildn force field parameters^67^ for the protein and TIP3P water model, along with ADP parameters generated by Carlson and accessed from the Amber Parameter Database^65–67^. The phosphorylated simulations also used custom phosphothreonine parameters described above^65,66^.

*Minimization and equilibration for the WT AMBER simulations.* Local energy minimization was performed in three separate steps in order to gradually introduce bond constraints. An OpenMM System was instantiated with no constraints on bonds or angles for the first minimization, which took place in vacuum (with crystallographic waters) with no constraints on bonds or angles. After this minimization, a new System was instantiated with constraints on the lengths of all bonds involving a hydrogen atom, and minimization was repeated. The structure and positions of all atoms were then put into a new OpenMM Modeller object, where TIP3P waters were added to a cubic box extending 11 Å beyond the outermost protein atoms, along with neutralizing counterions and sufficient excess NaCl to achieve an effective salt concentration of 300 mM. Another System with constrained bonds to hydrogen was created from the solvated structure and minimized. To minimally relax the structure before deploying simulations to Folding@Home, 5000 steps of Langevin dynamics were run using a Langevin integrator with a time step of 2.0 fs, temperature of 300.0 K, and collision rate of 5.0 ps^−1^. Nonbonded forces were modeled using the particle-mesh Ewald (PME) method with default parameters with a cutoff distance of 9.0 Å. All other settings remained at default values, except double precision was used throughout the minimization-and-equilibration process.

*Production simulation for WT amber simulations.* The resulting system, integrator, and state data from minimal equilibrations were serialized to XML format for simulation on Folding@Home using a simulation core based on OpenMM 6.3^61,68^ for both the phosphorylated and unphosphorylated systems. This entire process was repeated 5 times each phosphorylation simulations with and without Tpx2 to set up individual Folding@home RUNs, with each RUN representing a distinct initial configuration generated by the minimization-and-equilibration procedure. For each of the RUNs, 50 CLONEs with different initial random velocities and random seeds were simulated on Folding@home, where each clone ran for a maximum of 500 ns (250 million Langevin dynamics steps of 2 fs timestep with all-atom output frames saved every 125,000 steps using single precision and a Monte Carlo Barostat with pressure of 1 atm, temperature of 300 Kelvin, and barostat update interval of 50 steps), generating over 100 μs of aggregate simulation data for each of the WT conditions (with and without Tpx2).

*Modeling Spin Probe-labeled AurA.* Because AMBER parameters for spin probes were not widely available, we used the equivalently modern CHARMM36 generation of forcefields for MTSL-labeled AurA, simulated in complex with ADP and with and without both TPX2 and phosphorylation, for a total of four possible combinations per starting structure. Simulations were started from two different starting configurations: DFG-in (PDB ID: 1OL5) and DFG-out (5L8K^50^). For the crystal structure of 1OL5, Schrödinger's PrepWizard^69^ (release 2016-4) was used to model in Tpx2 residues 23-29 (unresolved in 1OL5), and add in hydrogens at pH 7.4 for both protein residues and ADP. The protonation state of ADP was assigned the lowest energy state using Epik at pH 7.4±2. Hydrogen bonding was optimized using PROPKA at pH 7.4±2. The entire structure was minimized using OPLS3 and an RMSD convergence cutoff of 0.3Å. Because TPX2 is not present in the 5L8K structure, the coordinates and crystal waters of Tpx2 in 1OL5 after preparation were transferred to the unprepared 5L8K after aligning the kinase domain to 1OL5 and deleting the vNAR domain (chain B) from 5L8K. 5L8K was then prepared using the same protocol as above, removing any organic solvent molecules but retaining all crystal waters. All three structures were run through CHARMM-GUI Solvator tool^70^ (http://www.charmm-gui.org/?doc=input/solvator). In the first stage of this tool, all crystal waters and magnesiums in the structures where retained, while sulfates were deleted. Tpx2 was deleted at this stage for the non-Tpx2 conditions. Phosphorylation was either built in or deleted for the threonine at residue 288 in the second stage of the Solvator tool. Also in this stage, residues 225 and 284 were mutated to cysteines and the MTSL spin label (named CYR1) was added to those residues. C290 was mutated to serine in the unphosphorylated conditions and alanine in the phosphorylated conditions, to match the DEER experimental conditions. After the PDB file was generated, a rectangular solvent box was generated using 10Å edge distance fit to the protein size, with 300 mM NaCl placed using the Monte-Carlo method.

*Parameterization the MTSL labeled CHARMM simulations.* An OpenMM ForceField was instantiated using CHARM36^71^ force field parameters for the protein and water model, along with ADP, TPO, and CYR1 parameter files output by CHARMM-GUI.

*Minimization and equilibration for the MTSL labeled CHARMM simulations.* A local energy minimization was performed with no constraints on bonds or angles, which took place in the solvated water box output by CHARMM-GUI and loaded into an OpenMM object. After minimization, 5000 steps of NVT dynamics were run using a Langevin integrator with a time step of 1.0 fs, temperature of 50K and a collision rate of 90.0 ps^−1^. Nonbonded forces were modeled using the particle-mesh Ewald (PME) method with a cutoff distance of 9.0 Å. All other settings remained at default values, except mixed precision was used throughout. After this, a second equilibration was run using 500000 steps of NPT dynamics using a Langevin integrator with a temperature of 300 Kelvin, collision rate of 90 ps^−1^, and timestep of 2.0 fs. A Monte Carlo Barostat was used with pressure of 1 atm and barostat update interval of 50 steps. To minimally relax this structure, 500000 steps of Langevin dynamics were run using a Langevin integrator with a 2 fs time step, 300 K temperature, and a collision rate of 5 ps^−1^.

*Production simulation for MTSL labeled CHARMM simulations.* The resulting system, integrator, and state data from the minimization and equilibration were serialized to XML format for simulation on Folding@Home using a simulation core based on OpenMM 6.3. This was done for all 8 conditions, where each combination of starting structure, phosphorylation status and Tpx2 status was set up as a RUN. For each of the RUNs, 100 CLONEs with different initial random velocities and random seeds were simulated on Folding@home, where each clone ran for a maximum of 3 μs (1.5 billion Langevin dynamics steps with all-atom output frames saved every 250,000 steps using mixed precision and a Monte Carlo Barostat with pressure of 1 atm, 300 Kelvin, and barostat frequency of 50). In aggregate, each of the 12 configurations totaled between 75-110μs per starting configuration.

*Data Analysis.* Distances and torsions were computed using the compute_distances and compute_dihedrals functions in MDTraj v 1.8.0. For the WT Amber simulations, the first 100 ns of each CLONE were discarded. The first 250 ns were discarded for the CHARMM MTSL labeled AurA simulations to allow sufficient relaxation following the introduction of spin probes. Distance probability plots were generated using Seaborn v0.8.1 (https://doi.org/10.5281/zenodo.883859) distplot using the norm_hist parameter. The distance and dihedral contour plots were generated using the kdeplot function in Seaborn v0.8.1. All analysis scripts have been made publically available^9^.

